# Cross-cancer pleiotropic associations with lung cancer risk in African Americans

**DOI:** 10.1101/405035

**Authors:** Carissa C. Jones, Yuki Bradford, Christopher I. Amos, William J. Blot, Stephen J. Chanock, Curtis C. Harris, Ann G. Schwartz, Margaret R. Spitz, John K. Wiencke, Margaret R. Wrensch, Xifeng Wu, Melinda C. Aldrich

**Affiliations:** Department of Thoracic Surgery, Vanderbilt University Medical Center, Nashville, TN; Vanderbilt Genetics Institute, Vanderbilt University Medical Center, Nashville, TN; School of Medicine, University of Pennsylvania, Philadelphia, PA; Department of Medicine, Baylor College of Medicine, Houston, TX; Division of Epidemiology, Department of Medicine, Vanderbilt University Medical Center, Nashville, TN; Division of Cancer Epidemiology and Genetics, National Cancer Institute, Bethesda, MD; Laboratory of Human Carcinogenesis, National Cancer Institute, Bethesda, MD; Karmanos Cancer Institute, Wayne State University, Detroit, MI; Department of Neurological Surgery, University of California San Francisco, San Francisco, CA; Department of Epidemiology and Biostatistics, University of California San Francisco, San Francisco, CA; Institute of Human Genetics, University of California San Francisco, San Francisco, CA; Department of Epidemiology, Division of Cancer Prevention and Population Sciences, University of Texas MD Anderson Cancer Center, Houston, TX; Department of Biomedical Informatics, Vanderbilt University Medical Center, Nashville, TN

## Abstract

**Background:** Identifying genetic variants with pleiotropic associations across multiple cancers can reveal shared biologic pathways. Prior pleiotropic studies have primarily focused on European descent individuals. Yet population-specific genetic variation can occur and potential pleiotropic associations among diverse racial/ethnic populations could be missed. We examined cross-cancer pleiotropic associations with lung cancer risk in African Americans.

**Methods:** We conducted a pleiotropic analysis among 1,410 African American lung cancer cases and 2,843 controls. We examined 36,958 variants previously associated (or in linkage disequilibrium) with cancer in prior genome-wide association studies. Logistic regression analyses were conducted, adjusting for age, sex, global ancestry, study site, and smoking status.

**Results:** We identified three novel genomic regions significantly associated (FDR-corrected p-value < 0.10) with lung cancer risk (rs336958 on 5q14.3, rs7186207 on 16q22.2, and rs11658063 on 17q12). On chromosome16q22.2, rs7186207 was significantly associated with increased risk (OR=1.24, 95% CI: 1.12-1.38) and functional annotation using GTEx showed rs7186207 modifies *DHODH* gene expression. The risk allele at rs336958 on 5q14.3 was associated with reduced lung cancer risk (OR=0.68, 95% CI: 0.56-0.82), while the risk allele at rs11658063 on 17q12 was associated with increased risk (OR=1.24, 95% CI: 1.11-1.39).

**Conclusion:** We identified novel associations on chromosomes 5q14.3, 16q22.2, and 17q12, which contain *HNF1B, DHODH,* and *HAPLN1* genes, respectively. SNPs within these regions have been previously associated with multiple cancers. This is the first study to examine cross-cancer pleiotropic associations for lung cancer in African Americans.

**Impact:** Our findings demonstrate novel cross-cancer pleiotropic associations with lung cancer risk in African Americans.

## Introduction

Pleiotropy occurs when a genetic locus is associated with more than one trait (1) and has been observed across multiple phenotypes, including cancer (2-5). These shared associations suggest potential common biologic pathways. One example of pleiotropy occurs at the *TERT* gene region. Germline mutations in *TERT* are associated with telomere length (6), but have also been associated with phenotypes, such as pulmonary fibrosis (7), red blood cell count (8), and aplastic anemia (9). *TERT* mutations have also been identified across multiple cancer types (10), including breast (11), prostate (12), pancreatic (13), glioma (14), and lung (15-17). Identifying pleiotropy is a useful tool for genetic association studies to discover common biologic mechanisms, shared genetic architecture across diseases, and potentially new opportunities for therapeutic targets.

Cross-cancer pleiotropic analysis has been conducted for several cancer types (2, 3, 5, 18), including lung cancer (4, 5). Pleiotropic analysis of lung cancer has identified associations with *LSP1, ADAM15/THBS3, CDKN2B-AS1*, and *BRCA2* genes in populations of predominantly European ancestry (4, 5). However, recent studies have shown the non-transferability of risk alleles across racial/ethnic groups (19, 20). It remains unknown whether pleiotropic associations of prior cancer variants are generalizable and associated with lung cancer among African American individuals. Furthermore, only a few studies (21) have accounted for either pleiotropy arising from variants in linkage disequilibrium or the ancestry of the discovery population. We used genome-wide genotyping data from five African American study populations to identify pleiotropic associations with lung cancer, incorporating variants in linkage disequilibrium and accounting for ancestry.

## Methods

### Study population

African American lung cancer cases and controls were selected from five study sites: the MD Anderson (MDA) Lung Cancer Epidemiology Study and Project CHURCH (Creating a Higher Understanding of Cancer Research & Community Health), NCI Lung Cancer Case-Control Study, the Northern California Lung Cancer Study from the University of California, San Francisco (UCSF), the Southern Community Cohort Study (SCCS), and three studies from the Karmanos Cancer Institute at Wayne State University (WSU), i.e. Family Health Study III; Women’s Epidemiology of Lung Disease Study; and Exploring Health, Ancestry and Lung Epidemiology Study. A detailed description of each study has been previously reported (22). All studies were approved by the Institutional Review Board at each institution and written informed consent was obtained from all participants.

### Genotyping, quality control, and imputation

Samples were previously genotyped (22) on the Illumina Human Hap 1M Duo array at the NCI Cancer Genomics Research Laboratory (CGR) in the Division of Cancer Epidemiology and Genetics (DCEG) at the National Cancer Institute. Genotyping data underwent strict quality control (Supplemental Figure 1). Briefly, single nucleotide polymorphisms (SNPs) were excluded if they were non-autosomal, had a MAF <1%, <95% genotyping efficiency, or a Hardy-Weinberg Equilibrium (HWE) p-value <1×10^−6^. Individuals were excluded if they had a <95% genotyping efficiency. Pairwise identity-by-descent was examined to identify related individuals; for each genetically related pair, the individual with the lowest genotyping efficiency was excluded (N=147). No individuals were excluded due to inconsistencies between reported and genetic sex, but four individuals with an “unknown” reported sex were filled in based on the calculated genetic sex. All quality control filtering was applied using PLINK (23).

Missing genotypes were imputed using IMPUTE2 (24), with pre-phasing performed using SHAPEIT (25-27). Haplotypes from the cosmopolitan 1000 Genomes phase 3 population consisting of 2,504 individuals from 26 countries were used as a reference population. SNPs imputed with low certainty were excluded based on an info score <0.4, MAF<0.01, and HWE p-value <1×10^−5^.

### Ancestry estimation

Supervised admixture analysis was performed using ADMIXTURE software (28) to obtain global estimates of African and European ancestry for all individuals. Admixture analysis was performed on observed genotypes merged with the CEU (CEPH Utah residents with Northern and Western European ancestry) and YRI (Yoruba from Ibadan, Nigeria) HapMap reference populations (29), and pruned to a set of unlinked variants (window size = 50, step size = 10, r^2^ > 0.1). A total of 140,591 variants remained for supervised (k=2) admixture analysis.

### Selection of variants for pleiotropic analysis

All variants previously associated with any cancer type were identified from the NHGRI-EBI GWAS catalog as of April 2016. Manual review excluded studies in which the outcome was not risk (e.g. survival, prognosis, toxicity, relapse). Studies assessing interactions were also excluded. SNPs from each of the remaining studies were aggregated into a single list, hereafter referred to as “reported SNPs.”

Because genotyping arrays are designed to capture genome-wide variation using as few SNPs as possible, the majority of reported associations are rarely causal, but rather correlated (or in linkage disequilibrium, LD) with the true causal variant (30, 31). Additionally, it is established that LD patterns differ between racial/ethnic groups (32). Since the vast majority of the GWAS-reported SNPs are identified in European- or Asian-descent populations, we further expanded our list of reported SNPs based on LD structure of the 1000 Genomes (phase 3) reference population (33, 34) similar to the race/ethnicity of the population reported in the GWAS Catalog study. For example, the CHB (Han Chinese in Beijing, China) reference population was used to identify SNPs in LD with variants reported in Asian-descent populations, while the CEU (CEPH Utah Residents with Northern and Western European Ancestry) reference population was used for SNPs reported in European-descent populations. For the admixed Latino/a and African American populations we used relevant continental reference populations: CEU, CHB, YRI (Yoruba in Ibadan, Nigeria), and MXL (Mexican Ancestry from Los Angeles) for Latino/a and CEU, YRI, and ASW (Americans of African Ancestry in SW United States) for African Americans. For each reported SNP, we extracted all SNPs with a r^2^>0.6 and +/-100kb of the reported SNP in the appropriate 1000 Genomes reference population(s) using PLINK pairwise LD estimation. A final list of SNPs for analysis was generated, hereafter referred to as the “selected SNPs.”

### Statistical analysis

Logistic regression was performed for each additively coded risk allele using SNPTEST to account for imputation probabilities (35). Age, sex, smoking status (current/former/never), global African ancestry, and study site were included as covariates in the logistic regression models. A Benjamini-Hochberg false discovery rate (FDR) correction was applied to p-values to account for multiple testing. Statistical significance was defined as an FDR-corrected p-value < 0.1. Exploratory strata-specific analyses were conducted by histologic subtype, smoking status (ever/never), and sex. A meta-analysis was also conducted summarizing results from each study (adjusting for age, sex, smoking status, and global African ancestry) using a fixed effect model in METAL (36).

## Results

### Descriptive Characteristics

A total of 4,253 African American individuals remained following quality control, with 1,410 lung cancer cases and 2,843 controls (Table 1). Forty-five percent of all individuals were male and the mean age at diagnosis was 58 years. Lung cancer cases were on average five years older than controls. Among cases, 55% of participants were current smokers and 37% were former smokers, while 43% of controls were never smokers. The median global African ancestry was similar among cases and controls (84% and 83%, respectively). The most frequent histologic cell type among cases was adenocarcinoma (45%), followed by squamous cell carcinoma (24%). Descriptive characteristics by study and case/control status are presented in Table 1.

**Table 1.** Descriptive characteristics of African American lung cancer cases and controls.

| Characteristic | MDA<br>N=1374 | NCI<br>N=583 | SCCS<br>N=506 | UCSF<br>N=986 | WSU<br>N=804 | Total |  |  |
| --- | --- | --- | --- | --- | --- | --- | --- | --- |
|  |  |  |  |  |  | Cases<br>N=1,410 | Controls<br>N=2,843 | Combined<br>N=4,253 |
| Status |  |  |  |  |  |  |  |  |
| Cases, N (%) | 373 (27.1) | 208 (35.7) | 168 (33.2) | 325 (33.0) | 336 (41.8) |  |  | 1,410 (33.2) |
| Controls, N (%) | 1001 (72.9) | 375 (64.3) | 338 (66.8) | 661 (67.0) | 468 (58.2) |  |  | 2,843 (66.8) |
| Sex |  |  |  |  |  |  |  |  |
| Male | 525 (38.2) | 308 (52.8) | 300 (59.3) | 450 (45.6) | 317 (39.4) | 698 (49.5) | 1,202 (42.3) | 1,900 (44.7) |
| Female | 849 (61.8) | 275 (47.2) | 206 (40.7) | 536 (54.4) | 487 (60.6) | 712 (50.5) | 1,641 (57.7) | 2,353 (55.3) |
| Mean age at diagnosis, yr (SD) | 52.2 (13.5) | 64.4 (9.6) | 55.9 (8.9) | 63.5 (11.2) | 60.3 (11.3) | 61.5 (10.5) | 56.9 (13.2) | 58.4 (12.6) |
| Smoking Status |  |  |  |  |  |  |  |  |
| Current, N (%) | 339 (24.8) | 196 (33.7) | 276 (54.8) | 364 (38.1) | 358 (44.6) | 778 (55.3) | 755 (27.0) | 1,533 (36.4) |
| Former, N (%) | 379 (27.7) | 250 (43.0) | 111 (22.0) | 361 (37.8) | 257 (32.0) | 520 (36.9) | 838 (29.9) | 1,358 (32.3) |
| Never, N (%) | 649 (47.5) | 135 (23.2) | 117 (23.2) | 230 (24.1) | 187 (23.3) | 110 (7.8) | 1,208 (43.1) | 1,318 (31.3) |
| Median African Ancestry, % | 82.6 | 82.4 | 88.3 | 81.9 | 82.9 | 83.6 | 83.1 | 83.2 |
| Histology |  |  |  |  |  |  |  |  |
| Adenocarcinoma, N (%) | 173 (46.4) | 97 (47.1) | 47 (31.3) | 145 (44.6) | 170 (50.6) | 632 (45.5) | - | 632 (45.5) |
| Squamous Cell, N (%) | 103 (27.6) | 53 (25.7) | 28 (18.7) | 82 (25.2) | 71 (21.1) | 337 (24.2) | - | 337 (24.2) |
| Large Cell, N (%) | 2 (0.5) | 5 (2.4) | 11 (7.3) | 4 (1.2) | 12 (3.6) | 34 (2.4) | - | 34 (2.4) |
| Small Cell, N (%) | 23 (6.2) | 2 (1.0) | 14 (9.3) | 20 (6.0) | 22 (6.8) | 81 (5.8) | - | 81 (5.8) |
| Other, N (%) | 72 (19.3) | 49 (23.8) | 50 (33.3) | 72 (22.2) | 63 (18.8) | 306 (22.0) | - | 306 (22.0) |

MDA=MD Anderson Cancer Center; NCI=National Cancer Institute; SCCS=Southern Community Cohort Study; UCSF=University
of California San Francisco; WSU=Wayne State University.

### Variant Selection

A total of 266 unique studies were extracted from the NHGRI-EBI GWAS Catalog based on the search term “neoplasm.” Forty-six studies were excluded after manual review (see Methods), resulting in 220 studies reporting associations for 959 unique SNPs (“reported SNPs”). Seventy-four percent (163 out of 220) of studies were conducted in European-descent populations, followed by 26% (57 of 220) in Asian-descent populations (Table 2). The admixed Latino/a and African American populations accounted for only 3% and 5% of prior GWAS cancer studies, respectively (Table 2). Of all reported SNPs, 629 were directly observed in the genotype data and an additional 294 were imputed. Thirty-six reported SNPs were not present in the 1000 Genomes reference populations used for LD-based selection of SNPs and were dropped from analysis. Application of the PLINK pairwise LD estimation method (r^2^>0.6 and +/-100kb) to all reported SNPs increased the number of SNPs from 959 to 39,010 (Table 2).

**Table 2.** Number of studies and variants reported in the NHGRI-EBI GWAS Catalog (as of April 2016) and the number of SNPs identified after LD-based selection.

| **Race/Ethnicity in GWAS Catalog** | **Number of Studies<sup>a</sup>** | **Number of SNPs** | **1000 Genomes Population Used for LD Selection** | **Number of SNPs after LD Selection** |
| --- | --- | --- | --- | --- |
| European | 163 | 743 | CEU | 29,727 |
| Asian | 57 | 217 | CHB | 10,625 |
| Latino | 6 | 20 | CEU, YRI, MXL, CHB | 968 |
| African American | 11 | 49 | CEU, YRI, ASW | 1,490 |
| **Total** | **220** | **959** |  | **39,010** |

<sup>a</sup>Total number of studies is less than the sum of populations because 11 studies included two or more racial/ethnic populations and are counted once for each population.

### Logistic Regression Analysis

Among the 39,010 selected SNPs, 1,772 were neither observed nor imputed in our African American population and 280 SNPs failed to meet post-imputation quality control filtering, resulting in 36,958 selected SNPs for analysis. Logistic regression analysis revealed 40 SNPs that were significantly associated with lung cancer risk (Figure 1 and Table 3). The most statistically significant association was identified on chromosome 15q25.1 for rs17486278 (per allele odds ratio (OR) = 1.41, 95% confidence interval (CI): 1.26-1.57, FDR-corrected p-value=2.32×10^−5^), followed by chromosome 5p15 (rs2853677, OR=0.79, 95% CI: 0.71-0.88, FDR-corrected p-value=0.04), Table 3). Three additional SNPs, rs336958 (5q14.3), rs7186207 (16q22.2), and rs11658063 (17q12) also had significant associations with lung cancer risk. The C allele at rs336958 on 5q14.3 was associated with reduced risk with an OR=0.68 and 95% CI: 0.56-0.82 (FDR-corrected p-value=0.06). The association on chromosome 16q22.2 consisted of four SNPs with similar effect sizes, though only one, rs7186207, surpassed a 10% FDR correction threshold (OR=1.24, 95% CI=1.12-1.38, FDR-corrected p-value = 0.04). On chromosome 17q12, the G allele at rs11658063 was associated with an increased risk of lung cancer (OR=1.24 and 95% CI: 1.11-1.39, FDR-corrected p-value=0.10). Similar results were observed when study sites were combined using fixed effect meta-analysis (data not shown).

**Figure 1.**
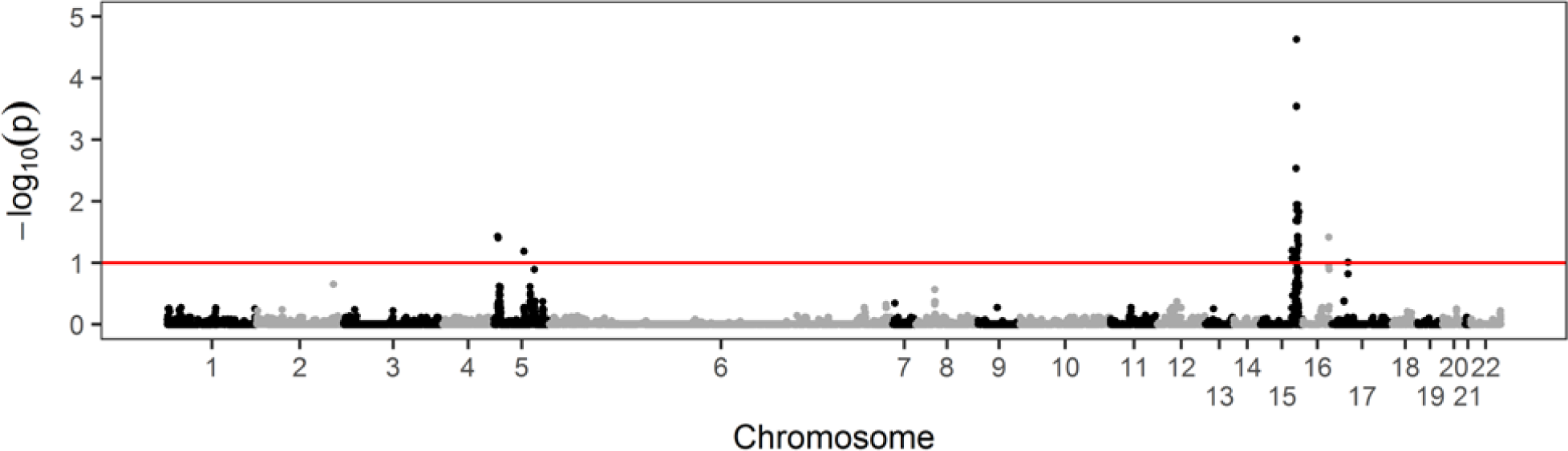
Logistic regression results for pleiotropic genetic associations with lung cancer risk among African American lung cancer cases and controls (N=4,253). Red line = 10% false discovery rate (FDR)

**Table 3.** SNPs significantly (FDR-corrected p-value < 0.10) associated with lung cancer risk among African American lung cancer cases and controls (N=4,253).

| SNP | Chr. | BP | Risk/<br>Ref.<br>Allele | Risk<br>Allele<br>Freq | Info<br>Score | OR | 95% CI | Unadjusted<br>P-value | FDR-corrected<br>P-value |
| --- | --- | --- | --- | --- | --- | --- | --- | --- | --- |
| rs17486278 | 15 | 78867482 | C/A | 0.3 | 1 | 1.41 | (1.26-1.57) | 6.27E-10 | 2.32x10<sup>-5</sup> |
| rs55781567 | 15 | 78857986 | G/C | 0.28 | 1 | 1.37 | (1.23-1.53) | 1.54E-08 | 2.84x10<sup>-4</sup> |
| rs2036527 | 15 | 78851615 | A/G | 0.23 | 1 | 1.36 | (1.21-1.54) | 2.87E-07 | 2.89x10<sup>-4</sup> |
| rs58365910 | 15 | 78849034 | C/T | 0.27 | 0.98 | 1.35 | (1.2-1.51) | 3.13E-07 | 2.89x10<sup>-4</sup> |
| rs147144681 | 15 | 78900908 | T/C | 0.18 | 1 | 1.36 | (1.2-1.55) | 2.14E-06 | 0.01 |
| rs576982 | 15 | 78870803 | T/C | 0.29 | 1 | 0.77 | (0.69-0.86) | 2.74E-06 | 0.01 |
| rs664172 | 15 | 78862762 | A/G | 0.28 | 1 | 0.76 | (0.68-0.85) | 2.17E-06 | 0.01 |
| rs667282 | 15 | 78863472 | C/T | 0.29 | 1 | 0.76 | (0.68-0.85) | 1.91E-06 | 0.01 |
| rs938682 | 15 | 78896547 | A/G | 0.72 | 1 | 1.3 | (1.17-1.46) | 2.56E-06 | 0.01 |
| rs569207 | 15 | 78873119 | T/C | 0.28 | 1 | 0.77 | (0.69-0.86) | 3.71E-06 | 0.01 |
| rs637137 | 15 | 78873976 | A/T | 0.29 | 1 | 0.77 | (0.69-0.86) | 4.15E-06 | 0.01 |
| rs11637630 | 15 | 78899719 | A/G | 0.71 | 0.99 | 1.29 | (1.16-1.44) | 5.35E-06 | 0.01 |
| rs2456020 | 15 | 78868398 | T/C | 0.4 | 1 | 0.79 | (0.71-0.87) | 5.52E-06 | 0.01 |
| rs55676755 | 15 | 78898932 | G/C | 0.17 | 1 | 1.35 | (1.19-1.54) | 5.69E-06 | 0.01 |
| rs7183604 | 15 | 78899213 | C/T | 0.72 | 0.99 | 1.3 | (1.16-1.45) | 4.87E-06 | 0.01 |
| rs12440014 | 15 | 78926726 | G/C | 0.22 | 0.91 | 0.75 | (0.66-0.85) | 6.39E-06 | 0.01 |
| rs3825845 | 15 | 78910258 | T/C | 0.23 | 1 | 0.76 | (0.68-0.86) | 8.24E-06 | 0.02 |
| rs503464 | 15 | 78857896 | A/T | 0.27 | 0.98 | 0.77 | (0.69-0.87) | 1.00E-05 | 0.02 |
| rs189218934 | 15 | 78903987 | C/T | 0.73 | 0.99 | 1.29 | (1.15-1.44) | 1.08E-05 | 0.02 |
| rs113931022 | 15 | 78901113 | T/C | 0.19 | 1 | 1.32 | (1.16-1.5) | 2.18E-05 | 0.04 |
| rs138544659 | 15 | 78900701 | G/T | 0.19 | 1 | 1.32 | (1.16-1.49) | 2.21E-05 | 0.04 |
| rs112878080 | 15 | 78900647 | G/A | 0.19 | 1 | 1.32 | (1.16-1.49) | 2.21E-05 | 0.04 |
| rs2853677 | 5 | 1287194 | A/G | 0.71 | 1 | 0.79 | (0.71-0.88) | 2.28E-05 | 0.04 |
| rs111704647 | 15 | 78900650 | T/C | 0.19 | 1 | 1.31 | (1.16-1.49) | 2.50E-05 | 0.04 |
| rs2735940 | 5 | 1296486 | G/A | 0.47 | 0.98 | 0.81 | (0.73-0.89) | 2.60E-05 | 0.04 |
| rs7186207 | 16 | 72035359 | C/T | 0.57 | 1 | 1.24 | (1.12-1.38) | 2.66E-05 | 0.04 |
| rs2853672 | 5 | 1292983 | A/C | 0.47 | 1 | 0.81 | (0.73-0.89) | 2.85E-05 | 0.04 |
| rs56077333 | 15 | 78899003 | A/C | 0.19 | 1 | 1.31 | (1.15-1.49) | 3.24E-05 | 0.04 |
| rs7170068 | 15 | 78912943 | A/G | 0.23 | 0.99 | 0.78 | (0.69-0.88) | 3.93E-05 | 0.05 |
| rs1051730 | 15 | 78894339 | A/G | 0.12 | 1 | 1.37 | (1.18-1.6) | 5.00E-05 | 0.06 |
| rs28491218 | 15 | 78267947 | C/T | 0.22 | 0.91 | 0.77 | (0.68-0.87) | 5.43E-05 | 0.06 |
| rs951266 | 15 | 78878541 | A/G | 0.11 | 0.99 | 1.39 | (1.18-1.63) | 5.28E-05 | 0.06 |
| rs12914385 | 15 | 78898723 | T/C | 0.2 | 1 | 1.29 | (1.14-1.46) | 5.69E-05 | 0.06 |
| rs336958 | 5 | 82973396 | C/T | 0.92 | 1 | 0.68 | (0.56-0.82) | 5.90E-05 | 0.06 |
| rs7172118 | 15 | 78862453 | A/C | 0.11 | 0.99 | 1.38 | (1.18-1.62) | 6.73E-05 | 0.07 |
| rs28360704 | 15 | 78268603 | T/C | 0.21 | 0.94 | 0.77 | (0.68-0.88) | 8.70E-05 | 0.08 |
| rs56390833 | 15 | 78877381 | A/C | 0.11 | 1 | 1.38 | (1.17-1.62) | 8.10E-05 | 0.08 |
| rs7180002 | 15 | 78873993 | T/A | 0.11 | 1 | 1.38 | (1.17-1.61) | 8.42E-05 | 0.08 |
| rs905739 | 15 | 78845110 | G/A | 0.25 | 0.98 | 0.79 | (0.7-0.89) | 8.77E-05 | 0.08 |
| rs11658063 | 17 | 36103872 | G/C | 0.61 | 0.88 | 1.24 | (1.11-1.39) | 1.04E-04 | 0.10 |
| rs16969968 | 15 | 78882925 | A/G | 0.07 | 1 | 1.49 | (1.22-1.83) | 1.14E-04 | 0.10 |
| rs8192482 | 15 | 78886198 | T/C | 0.07 | 1 | 1.49 | (1.22-1.83) | 1.18E-04 | 0.10 |

### Exploratory Stratified Analyses

To identify histologic subtype-specific associations for lung cancer risk, we examined adenocarcinoma cases (N=632) and squamous cell carcinoma cases (N=337) separately. No genetic variant was significantly associated with lung cancer risk in either histological subtype (Supplemental Figure 2). On chromosome 15q25.1, stratification by sex and smoking status revealed a sex-specific association among females (rs17486278, OR=1.51, 95% CI=1.30-1.76, FDR-corrected p-value=4.29×10^−3^, Supplemental Figure 3 and Supplemental Table 1) and among ever smokers (rs17486278, OR=1.41, 95% CI=1.26-1.58, FDR-corrected p-value=1.42×10^−4^, Supplemental Figure 4). One additional SNP, rs7486184 on chromosome 12q21.32 was also significantly associated with lung cancer risk among females (OR=1.35, 95% CI=1.17-1.55, FDR-corrected p-value=0.10, Supplemental Figure 3 and Supplemental Table 1). No SNPs were significant after an FDR correction in males (Supplemental Figure 3). Among ever smoking African Americans, 33 SNPs on chromosomes 5p15.33 and 16q22.2 had FDR-corrected p-values ≤ 0.10 (Supplemental Figure 4 and Supplemental Table 2). No p-values were statistically significant after FDR correction in never smokers (Supplemental Figure 4).

## Discussion

The present analysis sought to identify cross-cancer pleiotropic genetic associations for lung cancer risk in African Americans. The two most significant associations were on chromosome 15q25.1 and 5p15.33, both of which have been previously associated with lung cancer in African Americans (37-39) and recently validated in a non-independent African American consortium study (22) that included cases and controls utilized in the present study. We also identified three novel associations on chromosomes 5q14.3, 16q22.2, and 17q12. Chromosome 16q22.2 was also observed among ever smokers and an additional region on 12q21.32 was specific to women. Despite not meeting our threshold for statistical significance, there was suggestive evidence for an association of 5p15.33 and 15q25.1 among never smokers. These results are consistent with prior research indicating 5p15.33 is associated with lung cancer risk among never smokers (15, 40) and contribute to the ongoing debate as to whether 15q25.1 is directly associated with lung cancer or mediated by smoking (41-45).

Excluding established risk loci 5p15.33 and 15q25.1, the most significant association was for rs7186207 on chromosome 16q22.2 (FDR-corrected p-value = 0.04). All four SNPs in this region (rs7186207, rs8051239, rs7195958, and rs3213422) had similar odds ratios and were in strong LD (r^2^ > 0.68) with each other in both African (YRI and ASW) and European (CEU) 1000 Genomes reference populations (46). Given the high degree of correlation between variants, it is unsurprising that risk allele frequencies were similar among the four SNPs, ranging from 0.56 to 0.62. None of the four SNPs were among the reported SNPs extracted from the GWAS Catalog, but were selected because of their strong LD (r^2^ = 0.75-0.78) with rs12597458, a variant previously associated with prostate cancer risk (12). The 16q22.2 region has been previously associated with prostate cancer (47). SNPs rs7186207, rs8051239, and rs7195958 are intergenic and located between *PKD1L3* and *DHODH* and do not appear to be located at sites with regulatory potential based on histone modification marks (Supplemental Figure 5). However, GTEx data (48) reveals rs7186207 is significantly associated with *DHODH* gene expression in lung (p-value=2.1×10^−7^, Figure 2) and other tissues. The remaining SNP at this locus, rs3213422, is located within the first intron of *DHODH* (Supplemental Figure 5) and encodes a missense mutation, though SIFT (49) and PolyPhen-2 (50) both predict the mutation to be tolerated/benign. Given its close proximity to the exon boundary within *DHODH*, variation at rs3213422 could also affect exon splicing and GTEx data (48) reveals rs3213422 is a splice QTL for *DHODH* in aortic artery tissue (p-value=2.7×10^−6^). ENCODE data reveal H3K4me3 and H3K27ac markers surrounding rs3213422, indicative of active promoters and regulatory elements, as well as evidence for transcription factor binding (Supplemental Figure 5).

**Figure 2.**
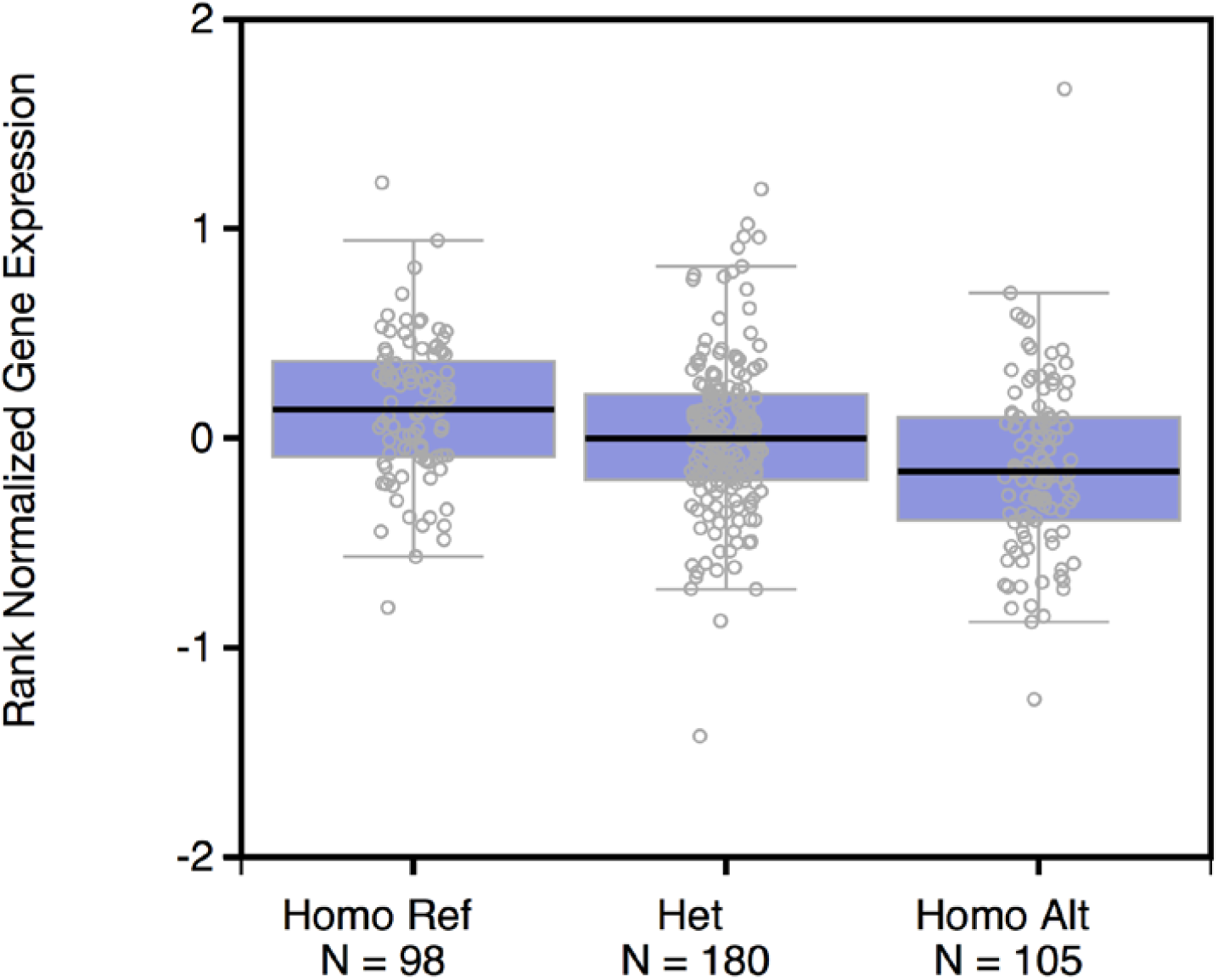
rs7186207 is an expression Quantitative Trait Loci (eQTL) for *DHODH* in lung tissue (p-value=2.1×10^−7^). Data are from GTEx Portal (48). Homo Ref=TT genotype; Het=TC genotype; Homo Alt=CC genotype

The *DHODH*, or dihydroorotate dehydrogenase, gene encodes a 43-kDa enzymatic protein localized to the inner mitochondrial membrane, where it interacts with the mitochondrial respiratory chain and acts as a rate-limiting step in *de novo* pyrimidine biosynthesis (51-53). Mutations within *DHODH* have been linked with Miller Syndrome, a recessive disorder characterized by malformations of the limbs and eyes, among other symptoms (54-57). *DHODH* has also been investigated for a role in cancer, including melanoma (58) and acute myeloid leukemia (59) and decreased expression of *DHODH* was associated with breast cancer risk (60). Several other studies have examined the utility of *DHODH* inhibitors in cancer by inducing cell cycle arrest and apoptosis in cancer cells (59, 61-66). While *DHODH* has not been previously associated with lung cancer risk, the abundance of biological evidence for its pleiotropic role in cancer gives credibility to the association.

We identified significant associations on chromosomes 5q14.3 and 17q12. Chromosome 5q14.3 SNP rs336958 is an intronic variant for *HAPLN1*, hyaluronan and proteoglycan link protein 1, which has been shown to play a role in cell adhesion and extracellular matrix structure. SNP rs336958 is in LD (r^2^=0.97) with rs4466137 which has been associated with prostate cancer risk (67). The larger 5q14.3 region has also been associated with prostate cancer (12), breast cancer (7), and Wilms tumors (68) and allelic imbalance in this region has been associated with multiple cancers, including lung (69-71).

The most significant SNP on chromosome 17q12 is rs11658063, a variant located in the first intron of *HNF1B*. HNF1 homeobox B (*HNF1B*) encodes a transcription factor and has been shown to play a role in cell development. SNPs in the *HNF1B* gene region have been previously associated with pancreatic (72), prostate (12, 73-79), ovarian (80-82), testicular (83), and endometrial (84-86) cancers. Expression of *HNF1B* has been associated with prognosis in hepatocellular carcinoma(87) and renal cell carcinoma (88). Furthermore, methylation of *HNF1B* has been observed in prostate (89), ovarian (82, 89-91), and lung (92) cancers and may have utility as a biomarker in ovarian cancer (90, 91).

The final notable region of association was on chromosome 12q21.32, where rs7486184 was associated with lung cancer in females. The intergenic variant rs7486184 is located approximately 40kb downstream of the *KITLG* gene and is in strong LD (r^2^=0.97) in Europeans (CEU) with reported variant rs995030 (46), which has been previously associated with testicular germ cell cancer in European-descent populations (93-95). Interestingly, rs7486184 and rs995030 may represent independent signals in African Americans since these SNPs are in weak LD in African descent populations (r^2^=0.15 for YRI and r^2^=0.42 for ASW) (46).

Of the SNPs previously reported to have a pleiotropic association with lung cancer (4, 5), two were not present in our current analysis. The remaining three SNPs (rs3817198, rs1057941, and rs4072037) were not significantly associated with lung cancer risk in our African American population. Both previous lung cancer pleiotropy studies (4, 5) utilized predominantly European-decent populations; thus, failure to replicate could represent population-specific effects. However, the minor allele frequency of rs3817198 and rs1057941 is higher in European versus African populations (rs3817198: 30% vs. 12%, respectively; rs1057941: 44% vs. 12%, respectively), which could result in reduced statistical power to detect these associations in our African American study population.

Previous pleiotropy studies have failed to consider differences in LD structure between racial/ethnic groups. Such considerations are important given recent publications noting the non-transferability of genetic risk predictions across diverse populations (19, 20). In the present analysis, a notable strength is our effort to expanded the list of reported SNPs by considering the LD structure of the racial/ethnic population of the discovery population, thus removing the assumption that the reported SNP has the same correlation structure, and therefore, tagging ability, with the causal SNP in all racial/ethnic groups. Importantly, it was through consideration of LD structure that the present study was able to identify novel lung cancer risk associations, as only two of the most significant SNPs (rs2853672 on 5p15.33 and rs1051730 on 15q25.1) were among the list of reported SNPs extracted from the GWAS catalog.

It is important to note that the present study is not independent of the study by Zanetti *et al*. as our cases and controls are a subset of the individuals in the Zanetti *et al* study. By restricting the analysis to SNPs with *a priori* evidence to examine cross-cancer pleiotropic associations, our study was able to identify novel lung cancer risk loci that may have been missed due to stringent multiple test corrections required in genome-wide association studies. Stratification by sex and smoking status revealed strong associations among women and ever smokers, suggesting the observed associations among all lung cancer cases and controls may be driven by these two subgroups. It remains to be determined whether the observed associations among women but not men represent a true biological phenomenon or are simply an artifact of reduced statistical power among men.

With our large sample size of African Americans and consistent results across the pooled and meta-analyses, we have identified several novel regions associated with lung cancer risk. Our findings highlight the need for a national effort dedicated to prioritizing research in diverse populations for future replication and fine mapping in African American lung cancer cases and controls to better understand the underlying mechanisms contributing to these pleiotropic signals. Functional studies should be performed to elucidate the pleiotropic effect of these associations across cancer types and identify common biological pathways across phenotypes that may lead to therapeutic targets for lung cancer.

## ACKNOWLEDGEMENTS

Data on SCCS cancer cases used in this publication were provided by the Alabama Statewide Cancer Registry; Kentucky Cancer Registry, Lexington, KY; Tennessee Department of Health, Office of Cancer Surveillance; Florida Cancer Data System; North Carolina Central Cancer Registry, North Carolina Division of Public Health; Georgia Comprehensive Cancer Registry; Louisiana Tumor Registry; Mississippi Cancer Registry; South Carolina Central Cancer Registry; Virginia Department of Health, Virginia Cancer Registry; Arkansas Department of Health, Cancer Registry, 4815 W. Markham, Little Rock, AR 72205. The Arkansas Central Cancer Registry is fully funded by a grant from National Program of Cancer Registries, Centers for Disease Control and Prevention (CDC). Data on SCCS cancer cases from Mississippi were collected by the Mississippi Cancer Registry, which participates in the National Program of Cancer Registries (NPCR) of the Centers for Disease Control and Prevention (CDC). The contents of this publication are solely the responsibility of the authors and do not necessarily represent the official views of the CDC or the Mississippi Cancer Registry.

The Genotype-Tissue Expression (GTEx) Project was supported by the Common Fund of the Office of the Director of the National Institutes of Health, and by NCI, NHGRI, NHLBI, NIDA, NIMH, and NINDS. The GTEx data used for the analyses described in this manuscript were obtained from the GTEx Portal on May 14, 2018.

## REFERENCES

1. Solovieff N, Cotsapas C, Lee PH, Purcell SM, Smoller JW. Pleiotropy in complex traits: challenges and strategies. Nature reviews Genetics. 2013;14:483–95.

2. Cheng I, Kocarnik JM, Dumitrescu L, Lindor NM, Chang-Claude J, Avery CL, et al. Pleiotropic effects of genetic risk variants for other cancers on colorectal cancer risk: PAGE, GECCO and CCFR consortia. Gut. 2014;63:800–7.

3. Panagiotou OA, Travis RC, Campa D, Berndt SI, Lindstrom S, Kraft P, et al. A genomewide pleiotropy scan for prostate cancer risk. European urology. 2014.

4. Park SL, Fesinmeyer MD, Timofeeva M, Caberto CP, Kocarnik JM, Han Y, et al. Pleiotropic associations of risk variants identified for other cancers with lung cancer risk: the PAGE and TRICL consortia. Journal of the National Cancer Institute. 2014;106:dju061.

5. Fehringer G, Kraft P, Pharoah PD, Eeles RA, Chatterjee N, Schumacher FR, et al. Cross-cancer genome-wide analysis of lung, ovary, breast, prostate, and colorectal cancer reveals novel pleiotropic associations. Cancer research. 2016;76:5103–14.

6. Liu Y, Cao L, Li Z, Zhou D, Liu W, Shen Q, et al. A genome-wide association study identifies a locus on TERT for mean telomere length in Han Chinese. PloS one. 2014;9:e85043.

7. Fingerlin TE, Murphy E, Zhang W, Peljto AL, Brown KK, Steele MP, et al. Genome-wide association study identifies multiple susceptibility loci for pulmonary fibrosis. Nature genetics. 2013;45:613–20.

8. Kamatani Y, Matsuda K, Okada Y, Kubo M, Hosono N, Daigo Y, et al. Genome-wide association study of hematological and biochemical traits in a Japanese population. Nature genetics. 2010;42:210–5.

9. Yamaguchi H, Calado RT, Ly H, Kajigaya S, Baerlocher GM, Chanock SJ, et al. Mutations in TERT, the gene for telomerase reverse transcriptase, in aplastic anemia. The New England journal of medicine. 2005;352:1413–24.

10. Mocellin S, Verdi D, Pooley KA, Landi MT, Egan KM, Baird DM, et al. Telomerase reverse transcriptase locus polymorphisms and cancer risk: a field synopsis and meta-analysis. Journal of the National Cancer Institute. 2012;104:840–54.

11. Couch FJ, Kuchenbaecker KB, Michailidou K, Mendoza-Fandino GA, Nord S, Lilyquist J, et al. Identification of four novel susceptibility loci for oestrogen receptor negative breast cancer. Nat Commun. 2016;7:11375.

12. Berndt SI, Wang Z, Yeager M, Alavanja MC, Albanes D, Amundadottir L, et al. Two susceptibility loci identified for prostate cancer aggressiveness. Nat Commun. 2015;6:6889.

13. Wolpin BM, Rizzato C, Kraft P, Kooperberg C, Petersen GM, Wang Z, et al. Genome-wide association study identifies multiple susceptibility loci for pancreatic cancer. Nature genetics. 2014;46:994–1000.

14. Kinnersley B, Labussiere M, Holroyd A, Di Stefano AL, Broderick P, Vijayakrishnan J, et al. Genome-wide association study identifies multiple susceptibility loci for glioma. Nat Commun. 2015;6:8559.

15. Hsiung CA, Lan Q, Hong YC, Chen CJ, Hosgood HD, Chang IS, et al. The 5p15.33 locus is associated with risk of lung adenocarcinoma in never-smoking females in Asia. PLoS genetics. 2010;6.

16. Landi MT, Chatterjee N, Yu K, Goldin LR, Goldstein AM, Rotunno M, et al. A genome-wide association study of lung cancer identifies a region of chromosome 5p15 associated with risk for adenocarcinoma. American journal of human genetics. 2009;85:679–91.

17. McKay JD, Hung RJ, Gaborieau V, Boffetta P, Chabrier A, Byrnes G, et al. Lung cancer susceptibility locus at 5p15.33. Nature genetics. 2008;40:1404–6.

18. Lindstrom S, Finucane H, Bulik-Sullivan B, Schumacher FR, Amos CI, Hung RJ, et al. Quantifying the genetic correlation between multiple cancer types. Cancer epidemiology, biomarkers & prevention: a publication of the American Association for Cancer Research, cosponsored by the American Society of Preventive Oncology. 2017;26:1427–35.

19. Manrai AK, Funke BH, Rehm HL, Olesen MS, Maron BA, Szolovits P, et al. Genetic misdiagnoses and the potential for health disparities. The New England journal of medicine. 2016;375:655–65.

20. Martin AR, Gignoux CR, Walters RK, Wojcik GL, Neale BM, Gravel S, et al. Human demographic history impacts genetic risk prediction across diverse populations. American journal of human genetics. 2017;100:635–49.

21. Wu YH, Graff RE, Passarelli MN, Hoffman JD, Ziv E, Hoffmann TJ, et al. Identification of Pleiotropic Cancer Susceptibility Variants from Genome-Wide Association Studies Reveals Functional Characteristics. Cancer epidemiology, biomarkers & prevention: a publication of the American Association for Cancer Research, cosponsored by the American Society of Preventive Oncology. 2018;27:75–85.

22. Zanetti KA, Wang Z, Aldrich M, Amos CI, Blot WJ, Bowman ED, et al. Genome-wide association study confirms lung cancer susceptibility loci on chromosomes 5p15 and 15q25 in an African-American population. Lung cancer. 2016;98:33–42.

23. Purcell S, Neale B, Todd-Brown K, Thomas L, Ferreira MA, Bender D, et al. PLINK: a tool set for whole-genome association and population-based linkage analyses. American journal of human genetics. 2007;81:559–75.

24. Howie BN, Donnelly P, Marchini J. A flexible and accurate genotype imputation method for the next generation of genome-wide association studies. PLoS genetics. 2009;5:e1000529.

25. Delaneau O, Coulonges C, Zagury JF. Shape-IT: new rapid and accurate algorithm for haplotype inference. BMC bioinformatics. 2008;9:540.

26. Delaneau O, Marchini J, Zagury JF. A linear complexity phasing method for thousands of genomes. Nature methods. 2011;9:179–81.

27. Delaneau O, Zagury JF, Marchini J. Improved whole-chromosome phasing for disease and population genetic studies. Nature methods. 2013;10:5–6.

28. Alexander DH, Novembre J, Lange K. Fast model-based estimation of ancestry in unrelated individuals. Genome research. 2009;19:1655–64.

29. International HapMap Consortium. The International HapMap Project. Nature. 2003;426:789–96.

30. Kruglyak L. Prospects for whole-genome linkage disequilibrium mapping of common disease genes. Nature genetics. 1999;22:139–44.

31. Ioannidis JP, Thomas G, Daly MJ. Validating, augmenting and refining genome-wide association signals. Nature reviews Genetics. 2009;10:318–29.

32. Evans DM, Cardon LR. A comparison of linkage disequilibrium patterns and estimated population recombination rates across multiple populations. American journal of human genetics. 2005;76:681–7.

33. Genomes Project C, Auton A, Brooks LD, Durbin RM, Garrison EP, Kang HM, et al. A global reference for human genetic variation. Nature. 2015;526:68–74.

34. Sudmant PH, Rausch T, Gardner EJ, Handsaker RE, Abyzov A, Huddleston J, et al. An integrated map of structural variation in 2,504 human genomes. Nature. 2015;526:75–81.

35. Marchini J, Howie B, Myers S, McVean G, Donnelly P. A new multipoint method for genome-wide association studies by imputation of genotypes. Nature genetics. 2007;39:906–13.

36. Willer CJ, Li Y, Abecasis GR. METAL: fast and efficient meta-analysis of genomewide association scans. Bioinformatics. 2010;26:2190–1.

37. Walsh KM, Amos CI, Wenzlaff AS, Gorlov IP, Sison JD, Wu X, et al. Association study of nicotinic acetylcholine receptor genes identifies a novel lung cancer susceptibility locus near CHRNA1 in African-Americans. Oncotarget. 2012;3:1428–38.

38. Walsh KM, Gorlov IP, Hansen HM, Wu X, Spitz MR, Zhang H, et al. Fine-mapping of the 5p15.33, 6p22.1-p21.31, and 15q25.1 regions identifies functional and histology-specific lung cancer susceptibility loci in African-Americans. Cancer epidemiology, biomarkers & prevention: a publication of the American Association for Cancer Research, cosponsored by the American Society of Preventive Oncology. 2013;22:251–60.

39. Schwartz AG, Cote ML, Wenzlaff AS, Land S, Amos CI. Racial differences in the association between SNPs on 15q25.1, smoking behavior, and risk of non-small cell lung cancer. Journal of thoracic oncology: official publication of the International Association for the Study of Lung Cancer. 2009;4:1195–201.

40. Wang Y, Broderick P, Matakidou A, Eisen T, Houlston RS. Role of 5p15.33 (TERT-CLPTM1L), 6p21.33 and 15q25.1 (CHRNA5-CHRNA3) variation and lung cancer risk in never-smokers. Carcinogenesis. 2010;31:234–8.

41. Hung RJ, McKay JD, Gaborieau V, Boffetta P, Hashibe M, Zaridze D, et al. A susceptibility locus for lung cancer maps to nicotinic acetylcholine receptor subunit genes on 15q25. Nature. 2008;452:633–7.

42. Thorgeirsson TE, Geller F, Sulem P, Rafnar T, Wiste A, Magnusson KP, et al. A variant associated with nicotine dependence, lung cancer and peripheral arterial disease. Nature. 2008;452:638–42.

43. Truong T, Hung RJ, Amos CI, Wu X, Bickeboller H, Rosenberger A, et al. Replication of lung cancer susceptibility loci at chromosomes 15q25, 5p15, and 6p21: a pooled analysis from the International Lung Cancer Consortium. Journal of the National Cancer Institute. 2010;102:959–71.

44. Wang Y, Broderick P, Matakidou A, Eisen T, Houlston RS. Chromosome 15q25 (CHRNA3-CHRNA5) variation impacts indirectly on lung cancer risk. PloS one. 2011;6:e19085.

45. Li Y, Sheu CC, Ye Y, de Andrade M, Wang L, Chang SC, et al. Genetic variants and risk of lung cancer in never smokers: a genome-wide association study. The Lancet Oncology. 2010;11:321–30.

46. Machiela MJ, Chanock SJ. LDlink: a web-based application for exploring population-specific haplotype structure and linking correlated alleles of possible functional variants. Bioinformatics. 2015;31:3555–7.

47. Al Olama AA, Kote-Jarai Z, Berndt SI, Conti DV, Schumacher F, Han Y, et al. A meta-analysis of 87,040 individuals identifies 23 new susceptibility loci for prostate cancer. Nature genetics. 2014;46:1103–9.

48. Consortium GT. The Genotype-Tissue Expression (GTEx) project. Nature genetics. 2013;45:580–5.

49. Kumar P, Henikoff S, Ng PC. Predicting the effects of coding non-synonymous variants on protein function using the SIFT algorithm. Nature protocols. 2009;4:1073–81.

50. Adzhubei IA, Schmidt S, Peshkin L, Ramensky VE, Gerasimova A, Bork P, et al. A method and server for predicting damaging missense mutations. Nature methods. 2010;7:248–9.

51. Evans DR, Guy HI. Mammalian pyrimidine biosynthesis: fresh insights into an ancient pathway. The Journal of biological chemistry. 2004;279:33035–8.

52. Fang J, Uchiumi T, Yagi M, Matsumoto S, Amamoto R, Takazaki S, et al. Dihydro-orotate dehydrogenase is physically associated with the respiratory complex and its loss leads to mitochondrial dysfunction. Biosci Rep. 2013;33:e00021.

53. Khutornenko AA, Roudko VV, Chernyak BV, Vartapetian AB, Chumakov PM, Evstafieva AG. Pyrimidine biosynthesis links mitochondrial respiration to the p53 pathway. Proceedings of the National Academy of Sciences of the United States of America. 2010;107:12828–33.

54. Fang J, Uchiumi T, Yagi M, Matsumoto S, Amamoto R, Saito T, et al. Protein instability and functional defects caused by mutations of dihydro-orotate dehydrogenase in Miller syndrome patients. Biosci Rep. 2012;32:631–9.

55. Kinoshita F, Kondoh T, Komori K, Matsui T, Harada N, Yanai A, et al. Miller syndrome with novel dihydroorotate dehydrogenase gene mutations. Pediatr Int. 2011;53:587–91.

56. Ng SB, Buckingham KJ, Lee C, Bigham AW, Tabor HK, Dent KM, et al. Exome sequencing identifies the cause of a mendelian disorder. Nature genetics. 2010;42:30–5.

57. Rainger J, Bengani H, Campbell L, Anderson E, Sokhi K, Lam W, et al. Miller (Genee-Wiedemann) syndrome represents a clinically and biochemically distinct subgroup of postaxial acrofacial dysostosis associated with partial deficiency of DHODH. Human molecular genetics. 2012;21:3969–83.

58. White RM, Cech J, Ratanasirintrawoot S, Lin CY, Rahl PB, Burke CJ, et al. DHODH modulates transcriptional elongation in the neural crest and melanoma. Nature. 2011;471:518–22.

59. Sykes DB, Kfoury YS, Mercier FE, Wawer MJ, Law JM, Haynes MK, et al. Inhibition of dihydroorotate dehydrogenase overcomes differentiation blockade in acute myeloid leukemia. Cell. 2016;167:171–86 e15.

60. Hoffman JD, Graff RE, Emami NC, Tai CG, Passarelli MN, Hu D, et al. Cis-eQTL-based trans-ethnic meta-analysis reveals novel genes associated with breast cancer risk. PLoS genetics. 2017;13:e1006690.

61. Baumann P, Mandl-Weber S, Volkl A, Adam C, Bumeder I, Oduncu F, et al. Dihydroorotate dehydrogenase inhibitor A771726 (leflunomide) induces apoptosis and diminishes proliferation of multiple myeloma cells. Molecular cancer therapeutics. 2009;8:366– 75.

62. He T, Haapa-Paananen S, Kaminskyy VO, Kohonen P, Fey V, Zhivotovsky B, et al. Inhibition of the mitochondrial pyrimidine biosynthesis enzyme dihydroorotate dehydrogenase by doxorubicin and brequinar sensitizes cancer cells to TRAIL-induced apoptosis. Oncogene. 2014;33:3538–49.

63. Zhu S, Yan X, Xiang Z, Ding HF, Cui H. Leflunomide reduces proliferation and induces apoptosis in neuroblastoma cells in vitro and in vivo. PloS one. 2013;8:e71555.

64. Liu L, Dong Z, Lei Q, Yang J, Hu H, Li Q, et al. Inactivation/deficiency of DHODH induces cell cycle arrest and programed cell death in melanoma. Oncotarget. 2017;8:112354–70.

65. Ladds M, van Leeuwen IMM, Drummond CJ, Chu S, Healy AR, Popova G, et al. A DHODH inhibitor increases p53 synthesis and enhances tumor cell killing by p53 degradation blockage. Nat Commun. 2018;9:1107.

66. Koundinya M, Sudhalter J, Courjaud A, Lionne B, Touyer G, Bonnet L, et al. Dependence on the Pyrimidine Biosynthetic Enzyme DHODH Is a Synthetic Lethal Vulnerability in Mutant KRAS-Driven Cancers. Cell Chem Biol. 2018.

67. Murabito JM, Rosenberg CL, Finger D, Kreger BE, Levy D, Splansky GL, et al. A genome-wide association study of breast and prostate cancer in the NHLBI’s Framingham Heart Study. BMC Med Genet. 2007;8 Suppl 1:S6.

68. Turnbull C, Perdeaux ER, Pernet D, Naranjo A, Renwick A, Seal S, et al. A genome-wide association study identifies susceptibility loci for Wilms tumor. Nature genetics. 2012;44:681–4.

69. Tavassoli M, Steingrimsdottir H, Pierce E, Jiang X, Alagoz M, Farzaneh F, et al. Loss of heterozygosity on chromosome 5q in ovarian cancer is frequently accompanied by TP53 mutation and identifies a tumour suppressor gene locus at 5q13.1-21. Br J Cancer. 1996;74:115– 9.

70. Achille A, Baron A, Zamboni G, Di Pace C, Orlandini S, Scarpa A. Chromosome 5 allelic losses are early events in tumours of the papilla of Vater and occur at sites similar to those of gastric cancer. Br J Cancer. 1998;78:1653–60.

71. Gorgoulis VG, Mariatos G, Manolis EN, Zacharatos P, Kotsinas A, Liloglou T, et al. Allelic imbalance at the 5q14 locus is associated with decreased apoptotic rate in non-small cell lung carcinomas (NSCLCs). Possible synergistic effect with p53 gene alterations on apoptosis. Lung cancer. 2000;28:211–24.

72. Klein AP, Wolpin BM, Risch HA, Stolzenberg-Solomon RZ, Mocci E, Zhang M, et al. Genome-wide meta-analysis identifies five new susceptibility loci for pancreatic cancer. Nat Commun. 2018;9:556.

73. Eeles RA, Kote-Jarai Z, Giles GG, Olama AA, Guy M, Jugurnauth SK, et al. Multiple newly identified loci associated with prostate cancer susceptibility. Nature genetics. 2008;40:316–21.

74. Thomas G, Jacobs KB, Yeager M, Kraft P, Wacholder S, Orr N, et al. Multiple loci identified in a genome-wide association study of prostate cancer. Nature genetics. 2008;40:310– 5.

75. Eeles RA, Kote-Jarai Z, Al Olama AA, Giles GG, Guy M, Severi G, et al. Identification of seven new prostate cancer susceptibility loci through a genome-wide association study. Nature genetics. 2009;41:1116–21.

76. Schumacher FR, Berndt SI, Siddiq A, Jacobs KB, Wang Z, Lindstrom S, et al. Genome-wide association study identifies new prostate cancer susceptibility loci. Human molecular genetics. 2011;20:3867–75.

77. Takata R, Akamatsu S, Kubo M, Takahashi A, Hosono N, Kawaguchi T, et al. Genome-wide association study identifies five new susceptibility loci for prostate cancer in the Japanese population. Nature genetics. 2010;42:751–4.

78. Lange EM, Johnson AM, Wang Y, Zuhlke KA, Lu Y, Ribado JV, et al. Genome-wide association scan for variants associated with early-onset prostate cancer. PloS one. 2014;9:e93436.

79. Gudmundsson J, Sulem P, Gudbjartsson DF, Blondal T, Gylfason A, Agnarsson BA, et al. Genome-wide association and replication studies identify four variants associated with prostate cancer susceptibility. Nature genetics. 2009;41:1122–6.

80. Kuchenbaecker KB, Ramus SJ, Tyrer J, Lee A, Shen HC, Beesley J, et al. Identification of six new susceptibility loci for invasive epithelial ovarian cancer. Nature genetics. 2015;47:164–71.

81. Pharoah PD, Tsai YY, Ramus SJ, Phelan CM, Goode EL, Lawrenson K, et al. GWAS meta-analysis and replication identifies three new susceptibility loci for ovarian cancer. Nature genetics. 2013;45:362-70, 70e1-2.

82. Shen H, Fridley BL, Song H, Lawrenson K, Cunningham JM, Ramus SJ, et al. Epigenetic analysis leads to identification of HNF1B as a subtype-specific susceptibility gene for ovarian cancer. Nat Commun. 2013;4:1628.

83. Kristiansen W, Karlsson R, Rounge TB, Whitington T, Andreassen BK, Magnusson PK, et al. Two new loci and gene sets related to sex determination and cancer progression are associated with susceptibility to testicular germ cell tumor. Human molecular genetics. 2015;24:4138–46.

84. Mandato VD, Farnetti E, Torricelli F, Abrate M, Casali B, Ciarlini G, et al. HNF1B polymorphism influences the prognosis of endometrial cancer patients: a cohort study. BMC Cancer. 2015;15:229.

85. Spurdle AB, Thompson DJ, Ahmed S, Ferguson K, Healey CS, O’Mara T, et al. Genome-wide association study identifies a common variant associated with risk of endometrial cancer. Nature genetics. 2011;43:451–4.

86. Setiawan VW, Haessler J, Schumacher F, Cote ML, Deelman E, Fesinmeyer MD, et al. HNF1B and endometrial cancer risk: results from the PAGE study. PloS one. 2012;7:e30390.

87. Yu DD, Jing YY, Guo SW, Ye F, Lu W, Li Q, et al. Overexpression of hepatocyte nuclear factor-1beta predicting poor prognosis is associated with biliary phenotype in patients with hepatocellular carcinoma. Sci Rep. 2015;5:13319.

88. Sun M, Tong P, Kong W, Dong B, Huang Y, Park IY, et al. HNF1B loss exacerbates the development of chromophobe renal cell carcinomas. Cancer research. 2017;77:5313–26.

89. Ross-Adams H, Ball S, Lawrenson K, Halim S, Russell R, Wells C, et al. HNF1B variants associate with promoter methylation and regulate gene networks activated in prostate and ovarian cancer. Oncotarget. 2016;7:74734–46.

90. Bubancova I, Kovarikova H, Laco J, Ruszova E, Dvorak O, Palicka V, et al. Next-generation sequencing approach in methylation analysis of HNF1B and GATA4 genes: searching for biomarkers in ovarian cancer. Int J Mol Sci. 2017;18.

91. Huang W, Cheng X, Ji J, Zhang J, Li Q. The application value of HNF-1beta transcription factor in the diagnosis of ovarian clear cell carcinoma. Int J Gynecol Pathol. 2016;35:66–71.

92. Shi YX, Wang Y, Li X, Zhang W, Zhou HH, Yin JY, et al. Genome-wide DNA methylation profiling reveals novel epigenetic signatures in squamous cell lung cancer. BMC Genomics. 2017;18:901.

93. Chung CC, Kanetsky PA, Wang Z, Hildebrandt MA, Koster R, Skotheim RI, et al. Meta-analysis identifies four new loci associated with testicular germ cell tumor. Nature genetics. 2013;45:680–5.

94. Rapley EA, Turnbull C, Al Olama AA, Dermitzakis ET, Linger R, Huddart RA, et al. A genome-wide association study of testicular germ cell tumor. Nature genetics. 2009;41:807–10.

95. Ruark E, Seal S, McDonald H, Zhang F, Elliot A, Lau K, et al. Identification of nine new susceptibility loci for testicular cancer, including variants near DAZL and PRDM14. Nature genetics. 2013;45:686–9.

